# Integrated organismal responses induced by ecologically-relevant *p*CO_2_ and temperature exposures in developing lake sturgeon

**DOI:** 10.1101/2023.09.28.559814

**Authors:** L. D. Belding, M. J. Thorstensen, A. R. Quijada-Rodriguez, W. S. Bugg, G. R. Yoon, A. R. Loeppky, G. J. P. Allen, A. N. Schoen, M. L. Earhart, C. Brandt, J. L. Ali, D. Weihrauch, K. M. Jeffries, W. G. Anderson

**Author notes:** Co-First Authors (L. D. Belding, M. J. Thorstensen).

## Abstract

Atmospheric CO_2_ and temperature are rising concurrently, and may have profound impacts on the transcriptional, physiological, and behavioral responses of aquatic organisms. Further, spring snow melt may cause transient increases of *p*CO_2_ in freshwater systems. Lake sturgeon (*Acipenser fulvescens*) groups were raised in current and projected levels of warming and *p*CO_2_. Following an overwintering period, lake sturgeon were exposed to a transient increase in *p*CO_2_, simulating a spring melt. Diverging transcriptional patterns were found in each group and metabolic rate was lower in the combined stressor group compared to others. Behavioral assays revealed no effect of environment on alarm cue responses or boldness, but there was a decrease in total activity following an acute CO_2_ exposure. These results demonstrate compensatory and compounding mechanisms of *p*CO_2_ and warming dependent on developmental conditions of a freshwater fish, and provide key information for responses to future climate change.

## Introduction

The impact of rising atmospheric CO_2_ on marine biota is undeniable, but the highly variable chemistry of freshwater ecosystems complicates our understanding of physiological responses to increased CO_2_ in freshwater environments^1–7^. Moreover, increases in atmospheric CO_2_ coincide with elevation in global temperatures^8,9^. This increase in CO_2_ can be especially prominent in flowing freshwater, which is often supersaturated with respect to the atmosphere^10,11^. The combined effect of increased temperature and *p*CO_2_ may induce cumulative or novel responses in aquatic ectotherms through stressor interactions not elicited by each factor individually^12^. These combined effects are likely to be profound throughout organismal development, but our understanding of the long-term impacts of acute exposures to elevated CO_2_ is still limited^13–15^. To better understand the effects of climate change in natural systems, it is necessary to determine how *p*CO_2_ and temperature act in concert over extended timescales throughout a freshwater organism’s life.

Early life is a critical period in an individual’s development, with environmental changes causing profound and potentially long-lasting effects on an organism’s capacity to respond to future environmental perturbations (i.e., developmental plasticity)^16^. Along with increasing atmospheric CO_2_, spring snowmelts can transiently increase *p*CO_2_ in freshwater, reaching 10,000 µatm of *p*CO_2_ in systems where it is consistently around 1,000 µatm^17^. These transient increases depend on local geochemical conditions and buffering capacity, such as in Boreal forest ecosystems^17–19^. As climate change will lead to fishes developing in environments with higher thermal and hypercapnic baselines, we reasoned that increased *p*CO_2_ and temperature exposure throughout early development could alter the capacity of aquatic organisms to tolerate a transient *p*CO_2_ challenge. Multiple response directions are reasonable in this circumstance^20^: either fish exposed to a more stressful developmental environment would be more tolerant to a later *p*CO_2_ challenge because they adusted their phenotypes during development, or the developmental environment decreased tolerance to the later challenge by reducing physiological capacity for responses^21,22^.

In this study, we used the lake sturgeon (*Acipenser fulvescens*), an ancient, long-lived fish with extensive phenotypic plasticity in responding to environmental stressors^23–27^. Because their capacity for plasticity is unusually high among vertebrates, lake sturgeon are valuable for studying mechanistic responses to a broad range of stressors at different magnitudes. The reasons for this plasticity are unknown, but large genome size may be a factor^28–30^. We studied the interacting effects of temperature and *p*CO_2_ throughout early development, followed by a transient increase of *p*CO_2_ to simulate a spring snowmelt near the end of their first year of life (Fig. 1). Specifically, immediately following fertilization, lake sturgeon eggs were incubated at 16°C and either 2,500 or 1,000 μatm *p*CO_2_. The 1,000 μatm *p*CO_2_ group was chosen as an ecologically-relevant exposure based on present *p*CO_2_ levels of ∼1,120 to ∼1,300 μatm in temperate freshwater lakes, and also represents current freshwater *p*CO_2_ levels in many systems^5,17,31^. The 2,500 μatm exposure was chosen based on the projected levels of *p*CO_2_ in freshwater ecosystems around the end of the 21^st^ century^5,32^. The 1,000 μatm *p*CO_2_ group was thus treated as a relative control against which the elevated *p*CO_2_ group was compared. Ten days post-hatch, a random subset of larvae from both *p*CO_2_ treatments were transferred to a second set of holding tanks and the temperature was slowly raised to 22°C, the highest temperature that wild lake sturgeon currently experience during summer months in the northern part of their range^24^. Larvae from all four treatments were then maintained in their respective treatments for the following five months. Then overwintering was simulated in all treatments by reducing temperature to 3°C for the following three months with food deprived for the final two months. Following overwintering, conditions of a spring snowmelt were simulated in which all treatments were exposed to an acute transient increase in *p*CO_2_ of 10,000 µatm for seven days, and temperature in all tanks was gradually increased to 16°C at a rate of 1°C·day^−1^. Once all tanks were at 16°C, *p*CO_2_ was decreased to developmental conditions of 2,500 or 1,000 µatm (Fig. 1).

**Figure 1.**
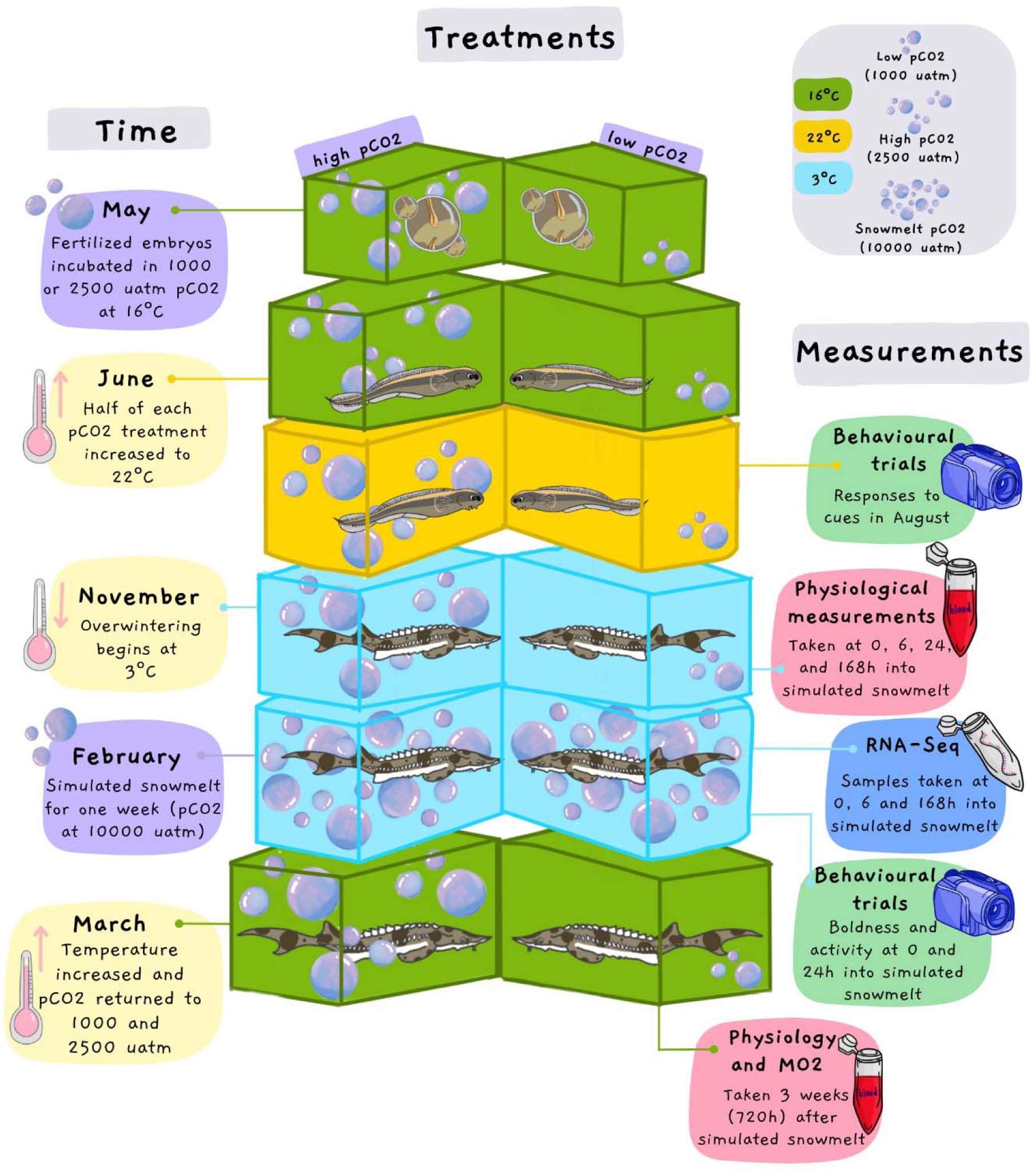
Conceptual diagram of rearing environments, transient *p*CO_2_ increase simulating a snowmelt, and sampling times. A timeline is provided on the left, and measurements taken at different times are shown on the right. Boxes correspond to discrete groups of fish, colours represent temperature at a given time for a group of fish while bubbles represent *p*CO_2_ in µatm.

After the transient acute *p*CO_2_ increase, lake sturgeon behaviour was analyzed for both boldness and alarm cue responses, in crossed rearing and exposure trial conditions. To test the effects of elevated long-term *p*CO_2_ and temperature exposures independently and together, changes in energy metabolism and acid-base regulation were examined using measurements of Na^+^/K^+^-ATPase activity, hematocrit, ammonia excretion, and metabolic rate at different timepoints following the transient *p*CO_2_ increase (Fig. 1). Messenger RNA (mRNA) sequencing was used to develop a mechanistic understanding of gill transcriptional responses, which was also assessed at different timepoints following the transient *p*CO_2_ increase (Fig. 1). A network analysis was performed among genes that changed over time within each experimental group to analyze regulatory processes versus those more proximate to biological responses. Therefore, this work connected long-term developmental conditions with a climate change-related stressor at multiple biological scales in a non-model fish, which will have implications for the mechanisms of climate change responses in freshwater organisms.

## Methods

### Animal Rearing

In the spring of 2017 gametes were collected from wild-caught spawning female and male lake sturgeon from the Winnipeg river, Manitoba and were transported to the University of Manitoba for fertilization. Approximately 50 ml of eggs from four females were mixed with approximately 100 ul of sperm from at least 10 different male sturgeon^53^. Following deadheasion, embryos were incubated at 16°C and either 2,500 or 1,000 μatm *p*CO_2_. Once the larvae hatched, bio-balls were added for substrate until exogenous feeding began. Prior to yolk-sac absoportion live artemia (Artemia International LLC, Texas, USA) were introduced to the tanks to encourage exogenous feeding. Two weeks following exogenous feeding, the lake sturgeon larvae diet was transitioned to bloodworms over approximately 30 days. Tanks were monitored three times daily, where any mortalities and debris were removed.

Ten days post-hatch, a random selection of larvae from both *p*CO_2_ treatments were transferred to a second set of holding tanks and the temperature was slowly raised to 22°C (1°C·day^−1^). Larvae from all four treatments (16°C 1,000 µatm (control), 16°C 2,500 µatm (high *p*CO_2_), 22°C 1,000 µatm (elevated temperature), and 22°C 2,500 µatm (high temperature and *p*CO_2_)) were held in these condtions for five months until they were exposed to overwintering conditions. Overwintering was simulated by decreasing the temperature in all treatments (1°C·day^−1^) until water temperature reached 3°C, following established protocols (Yoon et al., 2019b). The sturgeon were then held at 3°C for three months, with complete food restriction in the final two months to simulate Manitoba winters.

Eight months post-hatch sturgeon were exposed to a simulated acute spring snow melt, where all treatments were exposed to an acute transient increase in *p*CO_2_ of 10,000 µatm for seven days. Water temperatures in all tanks were then slowly increased to 16°C at a rate of 1°C·day^−1^. Once all tanks were at 16°C, *p*CO_2_ was decreased to developmental conditions of 2,500 or 1,000 µatm. Sampling for messenger RNA and physiology was conducted with respect to the start of the acute transient increase to 10,000 µatm of *p*CO_2_. Messenger RNA was collected at 0, 6, and 168 hours after the beginning of the increase. Na^+^/K^+^-ATPase activity and hematocrit was assessed at 0, 6, 24, 168, and 720 hours after it. Ammonia excretion was assessed 0, 24, 168, and 720 hours after. Routine and maximum metabolic rates were assessed 720 hours after the beginning of the increase. Time in alarm cue was measured 0 and 24 hours after. All animals used in this study were reared and sampled according to animal use and care guidelines established by the Canadian Council for Animal Care and approved by the Animal Care Committee at the University of Manitoba (protocol #F15-007).

### Messenger RNA Sequencing and Bioinformatics

Lake sturgeon sampled for this study were netted and euthanized by immersion in an overdose of tricaine methanesulfonate solution (250 mg L-1; MS-222, Syndel Laboratory, Vancouver, Canada). Gill tissue was then extracted from each fish and preserved in RNAlater (Invitrogen; Carlsbad, CA, USA), held at 4°C for 24 h and stored at −80°C prior to RNA extraction. Total RNA was extracted from the gill tissue by homogenization in 500 µl of lysis buffer (PureLink RNA Mini Kit; Invitrogen; Ambion Life Technologies; Waltham, MA, USA) for 10 min at 50 Hz using a TissueLyser II (Qiagen; Germantown, MD, USA) and then using a PureLink RNA Mini Kit (Invitrogen; Ambion Life Technologies) following manufacturer instructions. Total RNA purity and concentration was evaluated for all samples using a NanoDrop One (Thermo Fisher Scientific) as well as gel electrophoresis to assess RNA integrity. Samples were then stored at −80°C prior to preparation for sequencing. Samples were normalized to 50 ng·µL^−1^ and sequenced with paired-end, 100 base pair reads on a NovaSeq 6000 sequencing platform (Illumina) at the McGill Applied Genomics Innovation Core sequencing facility (https://www.mcgillgenomecentre.ca/). Prior to sequencing, samples were run on a Bioanalyzer to assess RNA quality and all samples had a minmum RNA Integrity Number of 8.0. Messenger RNA isolation was done with NEBNext Poly(A) Magnetic Isolation Modules (New England Biolabs) followed by the production of stranded cDNA libraries with NEBNext Ultra II Directional RNA Library Prep Kit for Illumina (New England Biolabs). Individual barcodes were applied with NEBNext dual adaptors (New England Biolabs) for each library prior to sequencing of 100 base pair paired-end reads.

A mean of 50.6 million raw reads were sequenced per individual (standard deviation 16.3 million raw reads) (Supplementary Information). The program fastp v0.20.1 was used for adapter trimming and read filtering, with a minimum phred quality of 15, a maximum of 40% of unqualified bases allowed in a read before read filtering, a minimum length of 100 base pairs, and polyG read tails were trimmed^54^. MultiQC v1.11 was used for collating sequencing and quality reports from fastp^55^. Salmon v1.7.0 was used to quantify trimmed and filtered raw reads against a lake sturgeon gill transcriptome^50,56^. The flags validateMappings, seqBias, and gcBias were included with the program to enable more selective alignments, correct for sequence-specific biases, and address potential GC biases, respectively.

The R package tximport v1.24.0 was used to import Salmon estimates of transcript abundance for differential gene expression analyses by edgeR v3.38.4^57–61^. The R package tidyverse v1.3.2 was useful throughout analyses in R^62^. Transcripts were not summarized to the level of gene models, because the polyploid lake sturgeon genome may have led to a fragmented transcriptome^50^. All differential expression analyses were thus performed at the transcript level. Transcripts were filtered for those present in any individual sample. A matrix of raw transcript counts was normalized to read length and effective library size following standard steps provided with tximport for using data with edgeR. A model design formula was used with no intercept and a combined treatment and time variable. Following posterior dispersion estimates with edgeR, a multidimensional scaling analysis was used to visualize variance among all individuals using the transcript abundances for the top 10% of transcripts by log_2_-fold change.

Because 66 potential pairwise contrasts were possible with the data (4 experimental groups at 3 timepoints), and yet more contrasts were possible with combinations of groups with respect to other combinations of groups, a strategy was needed to prioritize certain contrasts. Molecular and physiological mechanisms of responses to interacting *p*CO_2_ and temperature were most relevant to the present study. Therefore, pairwise contrasts were drawn between timepoints within groups: 0 versus 6 hours, 6 hours versus 168 hours, and 0 hours versus 168 hours. Significant transcripts with differential expression within these pairwise contrasts were collected by experimental group. Thus a list of transcripts with any change over time for each of the four experimental groups was generated. Transcripts that changed in abundance over time in the control conditions were subtracted from transcripts in the other groups, to identify transcripts that were specific to the environmental effects of increased temperature, *p*CO_2_, or both. In addition, we were interested in molecular mechanisms specific to each experimental group. We thus created a list of transcripts that changed in abundance over time within each group, unique to that group, by subtracting out both transcripts that changed over time in any other group. For example, transcripts unique to the interacting *p*CO_2_ and temperature were identified by subtracting out transcripts that changed between any timepoints in each of the control, *p*CO_2_, and temperature groups. A heatmap was created with counts per million values of these unique transcripts for *p*CO_2_, temperature, and *p*CO_2_ and temperature using pheatmap v1.0.12. For each pairwise contrast used, a genewise negative binomial generalized linear model with quasi-likelihood test was used (glmQLFit), where transcripts significant at Benjamini-Hochberg adjusted *p*-values (*q*)<0.05 were accepted as significant.

Significant transcripts were associated with gene annotations from the gill transcriptomes and used in gene set enrichment analyses with enrichR v3.1^63–65^. The enrichment databases Biological Process 2021 was used in each analysis. Gene ontology terms were accepted as significant at *q*<0.05, and considered in terms of combined scores, which is calculated using both *p*-values from a Fisher exact test and z-scores for deviations from expected rank^63^. For a gene ontology term that was of particular interest for its presence in both the specific and unique *p*CO_2_ and temperature enrichment analyses (negative regulation of organ growth; GO:0046621), gene function and counts per million among timepoints and experimental groups was further investigated.

In addition, a network analysis of genes annotated to transcripts that were specific to each treatment (temperature, *p*CO_2_, or both) but *not* unique was run with OmicsNet 2.0 using the zebrafish (*Danio rerio*) annotation database and GeneBankIDs^66,67^. Betweenness scores from genes within each network were assessed for skewness to quantify the distribution in centrality for genes in different networks using the moments v0.14.1 package in R. Genes were plotted in order of decreasing betweenness, omitting genes with betweenness of 0, with betweenness on the Y-axis to visualize the distribution of betweenness values in each experimental treatment.

EnrichR was also used with the Biological Process 2021 database to analyze gene ontology terms significant within network analyses for each treatment, where terms were prioritized by combined score. However, two enrichment analyses were run on each treatment: one on genes where betweenness was > 0, or the ‘hubs’ of the networks, and one on genes where betweenness = 0, or the ‘spokes’ of the networks. This partition in each treatment was done to distinguish between terms comprised of regulatory genes, versus others comprised of genes that were more proximate to biological processes.

### Na^+^/K^+^-ATPase Activity

Na^+^/K^+^-ATPase activity was assayed following protocols published in ^68^. A Bayesian approach was used to statistically assess Na^+^/K^+^-ATPase activity in each experimental group with the R package brms v2.18.0^69–71^. The model formula used was Na^+^/K^+^-ATPase activity dependent on experimental group as a fixed effect, with measurement run and individual specified as crossed random intercepts because individuals were measured in duplicate and measurement run may have introduced bias into the results. Variance was allowed to be unequal among experimental groups. Three priors were used: a normal prior of mean 0 and standard deviation 20 for the overall intercepts, a normal prior of mean 0 and standard deviation 20 for coefficients, and a Cauchy prior of mean 0 and standard deviation 4 for standard deviations.

Each model was run with a skew normal distribution over 5,000 Hamiltonian Monte Carlo warm-up iterations and 15,000 sampling iterations, with 4 separate chains used for sampling (60,000 sampling iterations total). Adapt delta was raised to 0.95 and tree depth was raised to 14 to address potential divergent transitions and maximum tree depth issues, respectively. Model fits were assessed with the potential scale reduction statistic RL (also called Rhat) and trace plots to estimate how well chain sampling and convergence progressed, as well as posterior predictive checks to visualize how well predicted results from the model matched real data. The model was accepted only if RL was 1.00 for all parameters.

Differences in Na^+^/K^+^-ATPase activity were assessed with 95% highest posterior density intervals using the R package emmeans v1.8.1-1, while Tidybayes v3.0.2 was used to visualize model results. Specifically, posterior distributions of Na^+^/K^+^-ATPase activity in each experimental group were compared in a pairwise manner with estimated marginal means to generate 95% highest posterior density intervals (HPD) of contrasts between groups. Estimated marginal means were also used to visualize posterior distributions for each experimental group. This modeling approach of fitting the dependent variable to experimental groups, choosing uninformative to weakly informative priors, controlling for variables when possible with random intercepts, checking models, using HPD intervals to make pairwise comparisons, and visualizing posterior distributions with estimated marginal means was applied to all models in the present study.

Correlations between Na^+^/K^+^-ATPase activity and mRNA transcript abundance were assessed with Bayesian models of acitivity dependent on transcript abundance, fit to gaussian distributions. Normal priors of mean 0 and standard deviation 100 were used for model coefficients and intercepts. One model was run for each of 33 transcripts annotated to subunits of the Na^+^/K^+^-ATPase gene. Expected log pointwise predictive densities were calculated for each model with the R package loo v2.6.0^72^.

### Hematocrit

Hematocrit measurements were done on lake sturgeon netted and euthanized as described above. To collect blood, a caudal severance of the tail was done, blood was collected in hematocrit tubes, and centrifuged in a micro-hematocrit centrifuge (CritSpin).

A Bayesian approach with brms was also used for assessing hematocrit amounts among different experimental groups. The model formula used was hematocrit dependent on the random slope of experimental group within timepoint, with variance allowed to be unequal using the same random effect formula. A skew normal distribution was used to fit the model, with a normal prior of mean 10 and standard deviation 30 for overall model intercept. Adapt delta was raised to 0.99 and tree depth raised to 12.

### Ammonia Excretion

For ammonia excretion measurements, lake sturgeon were placed in a 1L Tupperware container with 400 mL of aerated water corresponding to the fish’s respective treatment (pCO_2_/temperature). After a 60 min period to allow the fish to relax after transfer, two 1 mL water samples were collected at 0 and 1 hours. Ammonia in the collected water samples was measured spectrophotometrically by the colorimetric salicylate-hypochlorite assay described in ^73^. Ammonia excretion rate was then calculated based on the change in ammonia concentration between the 0 and 1 hour sample.

Ammonia excretion was assessed with a Bayesian model in brms, with ammonia excretion dependent on experimental group, and variance allowed to be unequal among experimental groups. A skew normal distribution was used to fit the model, with a normal prior of mean 0 and standard deviation 1 for model estimates and the overall intercept.

### Metabolic Rate at 720h Post-pCO_2_ Exposure

Metabolic rate was indirectly assessed by measuring whole-body oxygen consumption rate with intermittent flow respirometry (Loligo Systems, Viborg, Denmark). The protocol of measuring metabolic rate was performed as previously described^74^. It is worth to note that logistic constraints led to an invalid measurement of metabolic rate during the time of snow melt experiment due to the insurmountable measurement noise from low respiration and high bodymass:volume. Thus, we chose to use data only at the 720 h timepoint. The respirometry consisted of eight acrylic chamabers (639.0 ± 15.5 mlW; mean ± standard deviation.) and non-oxygen permeable tubing (63.3 ± 13.3 ml) submerged in a 200 L oval-shaped tank. Each chamber was comprised of flush tubing and also recirculation tubing with a flow-through chamber in which an oxygen dipping probe was placed to measure oxygen levels of air saturation at 1 Hz by Witrox 4 Oxygen Meter (Loligo Systems, Viborg, Denmark). Fish were fasted overnight prior to the measurement of metabolic rate. Then, fish were haphazardly captured from the rearing tank and placed them into the metabolic chambers. Black curtains were hung around the setup to minimize the visual disturbance. Fish were left in the metabolic chambers for 24 hours, after which fish were removed from the chambers and chased by gentle prodding to the tail for 15 minutes. Fish were immediately returned to the chambers and measured for two additional measurement cycles. Before and after each trial, oxygen consumption was measured without fish to estimate background respiration, and two data points were linearly interpolated over time and all the metabolic rate data were corrected (Rodgers et al. 2016). Only slopes with *R*^2^>0.9 were used for the data analysis. Routine metabolic rate (RMR) was calculated by averaging the metabolic rate data following the exlucision of first six hours of data as an acclimation. Maximum metabolic rate (MMR) was the highest data following the chase protocol. The average ratio between body mass and chamber volume was 81.8 ± 26.1, and the average ratio between background respiration and metabolic rate was 0.17 ± 0.29.

Bayesian models were used with either routine or maximum metabolic rate dependent on experimental group, a random intercept of date the measurement was taken, and variance was allowed to differ among experimental groups. Adapt delta was raised to 0.99. In addition, for specific contrasts of interest in metabolic rate model results, evidence ratios and posterior probabilities of differences were calculated with the *hypothesis* function in brms.

### Alarm Cue Responses at 0 and 24h Post-pCO_2_ Exposure

Fish were exposed to an alarm cue in a double-flume experiment immediately after exposure to 10,000 µatm of *p*CO_2_ and 24h post-exposure, in all 4 rearing groups (control, temperature, *p*CO_2_, and *p*CO_2_+temperature). Fish were recorded by video during the experiment. Proportion of time fish spent in the alarm cue was assessed in a blinded manner with experimental treatment and timepoint unknown to one assessor *via* randomized video identification numbers. To model the proportion of time that fish spent in the alarm cue, treatment was modeled as a random slope with timepoint as a random intercept, nested random intercepts of treatment tank ID within cue application side, and the date the experiment was performed as a random intercept. The model formula was thus: Proportion of time in cue ∼ (Treatment | Time) + (1 | Tank / Side) + (1 | Date). Variance was allowed to differ among a combined variable of treatment and timepoints. A normal prior of mean 50 and standard deviation 40 was used for the overall model intercept. Adapt delta raised to 0.99 and maximum tree depth raised to 12.

### Boldness and Activity in Crossed Rearing and Trial Groups

To assess the effects of rearing environment on behavioural responses to future increases in temperature, *p*CO_2_, or both, an experiment with crossed rearing and trial exposures was performed with a novel stimulus between 50 and 53 days post hatch. The novel stimulus was a plastic object previously unencountered by study subjects. Each cue of a control, alarm, food, or alarm and food was added to the tank and mixed around the object based on novel object work in ^1^. Fish from one of the four rearing groups was then added into the thigmotaxis (outer) zone of the tank. The experimenter then left the testing area within 20 seconds of adding the cue. Recording proceeded for 5 minutes, while data were only recorded and analyzed from the middle 3 minutes of video to reduce disturbances or behavioural abnormalities at the beginning or end of each trial. Time near the novel object, time in the thigmotaxis zone, total activity in any area of the tank, activity near the object, and activity in the thigmotaxis zone were measured in blinded recordings *via* randomized video identification numbers.

For time near the novel object, a skew normal distribution was used with time near the novel object dependent on a combined variable of rearing and trial group modeled with a random slope with cue as a random intercept, treatment tank nested within rearing tank as a random intercept, and the interaction of length and mass as a fixed effect. A normal prior of mean 0 and standard deviation 20 was used for the overall model intercept and fixed effect estimates, while a half-cauchy prior with scale parameter of 10 was used for model standard deviations. Adapt delta raised to 0.99 and maximum tree depth raised to 14. Time in thigmotaxis zone was run with identical model parameters, except with a student t-distribution used to fit the model overall.

Total activity in any area of the tank was modeled with a skew normal distribution, with total activity dependent on the same variables as the model for time near the novel object. A normal prior of mean 20 and standard deviation 50 was used for the overall model intercept and fixed effect estimates, while a half-cauchy prior with scale parameter 5 was used for model standard deviations. Activity near the novel object was modeled with the same independent variables, but we could not model variance as unequal among rearing and trial group combinations without numerous divergent transitions. As such, variance was modeled as equal among rearing and trial group combinations, as is typical for linear regression. Here, a normal prior of mean 0 and standard deviation 10 was used for the overall model intercept and fixed effect estimates, while a half-cauchy prior with scale parameter 5 was used for model standard deviations. Last, activity in the thigmotaxis zone was modeled with the same structure as time near the novel object. Here, a normal prior of mean 0 and standard deviation 20 was used for the overall model intercept and fixed effect estimates, while a half-cauchy prior with scale parameter 5 was used for model standard deviations.

## Results

### Transcriptomics

The top 10% of the transcripts (16,256) by log_2_-fold change were used in a multidimensional scaling analysis, which revealed variation consistent with *p*CO_2_ and temperature separately on dimension 1, and variation largely driven by time on dimension 2 (Fig. 2; Supplementary Fig. S1). Within gene ontology terms specific, but not unique, to the treatment groups, negative regulation of organ growth (GO:0046621) had the highest combined score (score) of 921,306 for the combined *p*CO_2_ and warm temperature treatment. The next highest term (regulation of organ growth; GO:0046620) had a score of 380 (Supplementary Information). Ear development (GO:0043583) was the third-highest term present in the combined *p*CO_2_ and temperature treatment with a score of 316, which was notable given potential connections of the stressors with otolith formation^32^. The *p*CO_2_-specific terms also showed a pattern of one gene ontology term with a higher score compared to others, with plus-end-directed vesicle transport along microtubule (GO:0072383) assigned a score of 902,620 while the next highest term had a score of 438 (Supplementary Information). The top two temperature-specific terms by combined score were also high compared to the rest, with lysosomal microautophagy (GO:0016237) and piecemeal microautophagy of the nucleus (GO:0034727) each assigned a score of 1,669,824 (Supplementary Information). Both of these terms shared the same set of genes (RB1CC1; ATG2A; ULK2; ULK1; ATG13; ATG7; ATG2B), and transcripts of all 7 were differentially expressed between timepoints within the warm temperature treatment and not in the control group.

**Figure 2.**
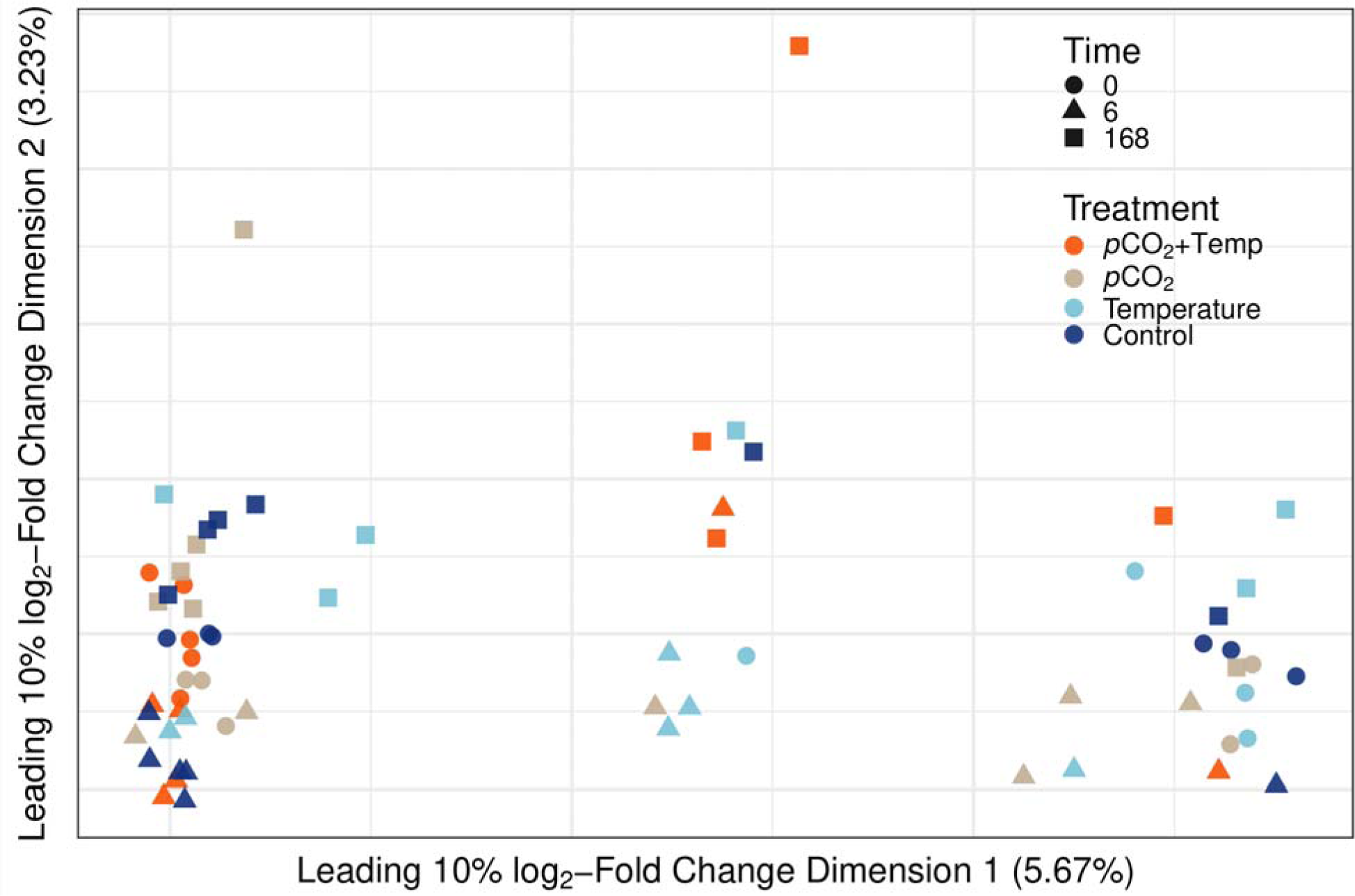
Multidimensional scaling plot of the top 10% of transcripts (16,256) by log_2_-fold change in the present experiment, with shapes representing time and colours representing experimental group. Messenger RNA data were collected 0, 6, and 168 hours into a transient *p*CO_2_ exposure of 10,000 µatm for 168 hours, simulating a spring snowmelt following the fish’s first year of life in the respective rearing environments and a simulated overwintering event.

Within gene ontology terms for transcripts unique to each treatment, negative regulation of organ growth (GO:0046621) also had the highest score of 616 for the combined *p*CO_2_ and temperature treatment, consistent with the overall treatment-specific results (Supplementary Information). A notable term unique to the *p*CO2 treatment was ear development (GO:0043583), with a score of 144 and was also present in overall terms in the combined *p*CO_2_ and temperature treatment. Interestingly, negative regulation of organ growth (GO:0046621) had the highest score in terms both specific and unique to the combined *p*CO_2_ and temperature group. Therefore, we investigated the genes present in this term and the patterns of mRNA transcript abundance more closely.

Five genes comprise the term negative regulation of organ growth: WWC1; WWC2; PTEN; WWC3; SLC6A4. Transcripts of all five genes were present in the combined *p*CO_2_ and temperature-specific terms, while *pten* was missing from the combined *p*CO_2_ and temperature-unique terms. Of the genes present in both the specific and unique analyses, the WWC genes are all a part of signaling pathways, involved in tumor suppression *via* restricting cell proliferation and promoting apoptosis. Meanwhile, SLC6A4 is sodium-dependent serotonin transporter, that terminates the action of and recycles serotonin, thus regulating serotonergic signaling (Uniprot). We investigated the mRNA transcript abundance associated with *slc6a4* in the data (transcript ID TR49140|c2_g1_i4). This transcript had a log_2_-fold change of −0.46 between the 168 h and 0 h timepoints (following acute *p*CO_2_ exposure) in the combined *p*CO_2_ and temperature treatment (*q*=0.011). We found a decrease of *slc6a4* transcript abundance in the combined *p*CO_2_ and temperature treatment, and limited differences in the other groups (Supplementary Fig. S2).

Temperature exhibited the most diffuse response in terms of network centrality, while *p*CO_2_ and temperature showed a response more focused on a few central genes (Fig. 3). Similar terms were present when analyzing genes more central in the networks of all three treatments. For example, the term SRP-dependent cotranslational protein targeting to membrane (GO:0006614) had the highest score in each of the isolated *p*CO_2_ and temperature treatments (27,487 and 11,298, respectively), and the second highest score in the combined *p*CO_2_ and temperature treatment (16,191). However, less consistent processes were apparent when analyzing ontology terms from genes on the edge of each network, which we reasoned may be more proximate to biological outcomes of regulatory pathways. Terms from the *p*CO_2_ treatment were related to splicing (e.g., spliceosomal conformational changes to generate catalytic conformation; GO:0000393; score 1,715,914) or protein processing (e.g., proteasomal ubiquitin-independent protein catabolic process; GO:0010499; score 871) (Supplementary Information).

**Figure 3.**
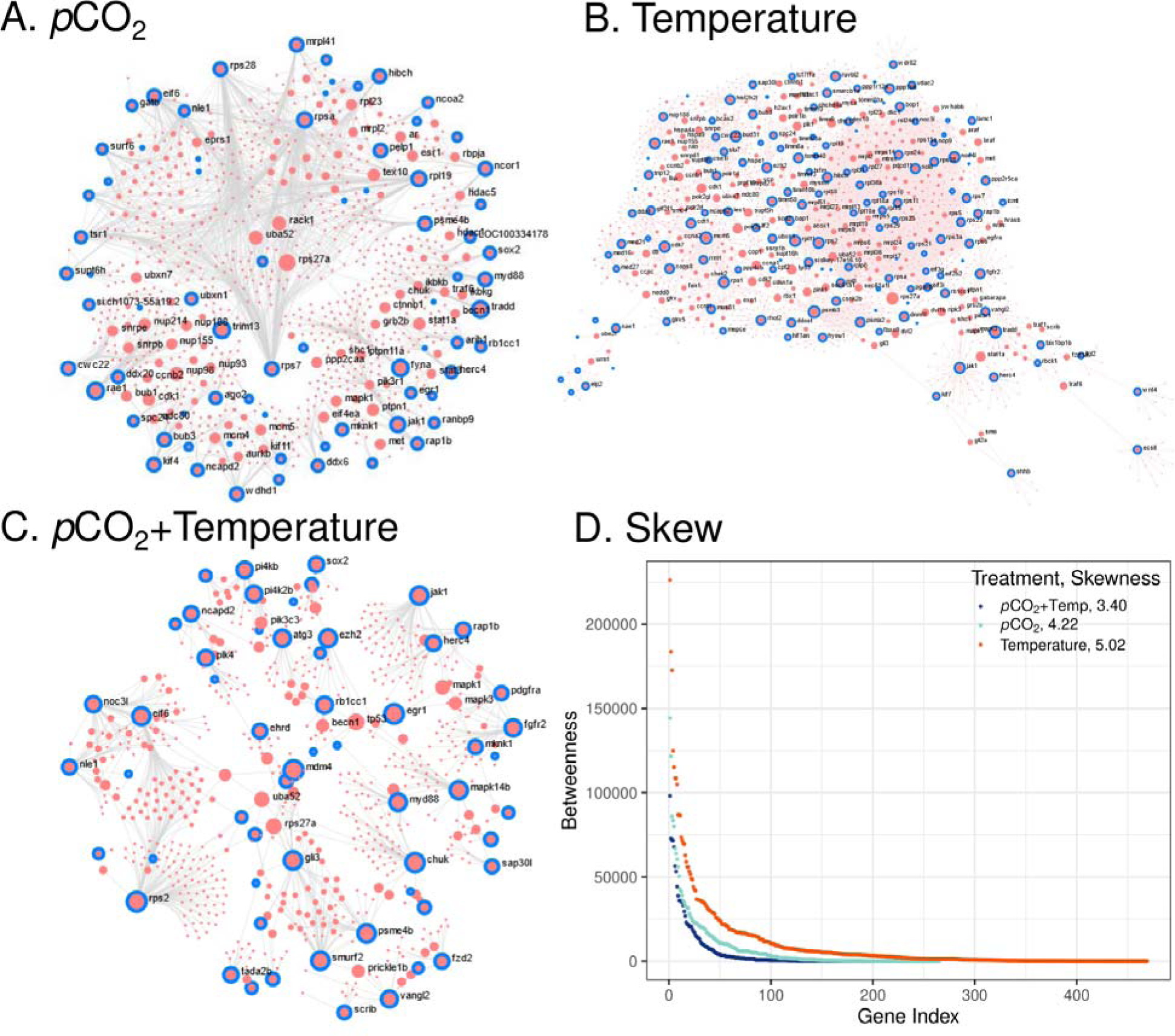
Transcriptional networks from genes with differential expression specific to each group following a simulated snowmelt and transient *p*CO_2_ increase of 10,000 µatm. Gene IDs were analyzed using OmicsNet 2.0 against the zebrafish (*Danio rerio*) protein interaction database. Panel A represents genes specific to fish reared in elevated *p*CO_2_ only, panel B represents genes specific to fish reared in elevated temperature only, and panel C represents genes specific to fish reared in both elevated *p*CO_2_ and temperature. Panel D shows ranked betweenness scores for genes of each group-specific network, with skewness values provided for the betweenness score distribution from each network.

Meanwhile, genes on the edge of the temperature treatment network were characterized by metabolism (e.g., regulation of cellular amine metabolic process; GO:0033238; score 1,438) or cell cycling (e.g., negative regulation of cell cycle G2/M phase transition; GO:1902750; score 1,385) (Supplementary Information). In the combined *p*CO_2_ and temperature treatment, genes along the edges of the network were characterized by translation regulation (e.g., cytoplasmic translation; GO:0002181; score 851) or cell death-related processes (e.g., positive regulation of intrinsic apoptotic signaling pathway by p53 class mediator; GO:1902255; score 540) (Supplementary Information). Therefore, a shared transcriptional response may have driven the networks initiated by all three experimental manipulations. However, *p*CO_2_ responses may have led to alternative splicing and protein ubiquitination, temperature responses to cell cycling and metabolism, and the combination of *p*CO_2_ and temperature to protein synthesis and cell death.

### Physiology

Na^+^/K^+^-ATPase activity increased between 0 and 168 hours in both the temperature and control groups, before returning to 0-hour levels at 720 hours post-acute *p*CO_2_ exposure (Fig. 4). Na^+^/K^+^-ATPase activity did not notably change in the *p*CO_2_ or combined *p*CO_2_ and temperature groups. There was no apparent trend in hematocrit in the experimental treatments, except it was lower in the control group at 0 hours than at 720 hours (Supplementary Fig. S3). Ammonia excretion was highest in the control group at 0 hours, but increased in all groups at 720 hours, especially the *p*CO_2_-only group (Supplementary Fig. S4). At 720 h, routine metabolic rate was lower in the combined *p*CO_2_ and temperature treatment than in the individual *p*CO_2_ and temperature treatments, a condition potentially linked to respiratory alkalosis (Supplementary Fig. S5)^33^. Similarly, maximum metabolic rate was slightly higher in the *p*CO_2_-only group relative to the combined *p*CO_2_ and temperature group (Supplementary Fig. S5).

**Figure 4.**
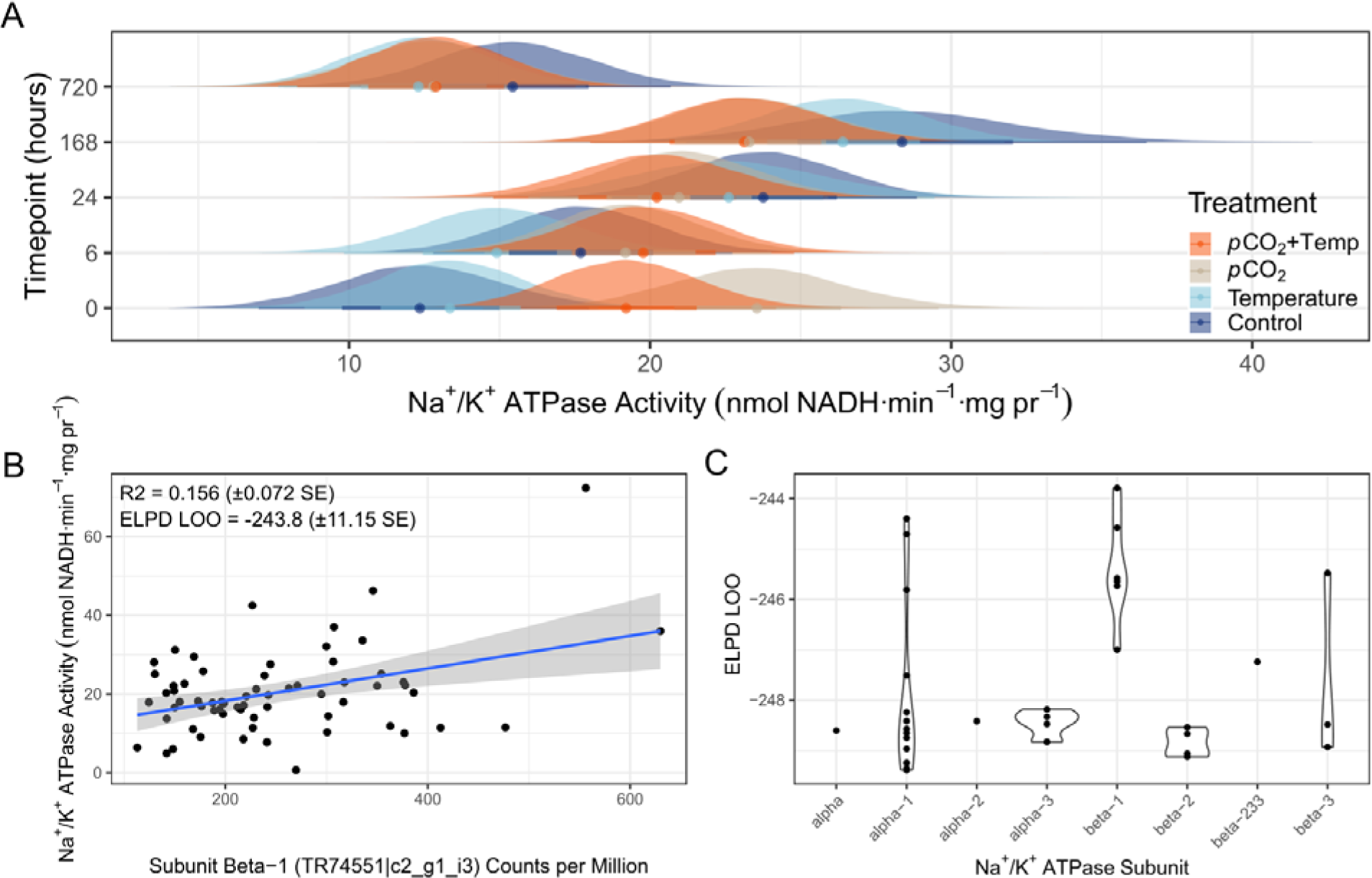
Posterior distributions, linear models, and expected log pointwise predictive densities calculated with leave one out cross validation (ELPD LOO) for Na^+^/K^+^-ATPase activity and associated transcript abundance. Panel A represents Bayesian posterior estimates of Na^+^/K^+^-ATPase activity calculated for each group, at each timepoint. Panel B represents the model of Na^+^/K^+^-ATPase activity dependent on Na^+^/K^+^-ATPase mRNA transcript abundance wit the lowest ELPD LOO value and highest *R*^2^ value out of 33 models of mRNA transcript abundance tested. Panel C shows ELPD LOO values for all 33 models of Na^+^/K^+^-ATPase mRNA transcript abundance correlated with enzyme activity, with Na^+^/K^+^-ATPase subunit annotations on the horizontal axis.

We correlated abundance of transcripts annotated to subunits of the gene Na^+^/K^+^-ATPase to enzyme activity. A total of 33 unique transcripts were annotated to subunits of the gene, with 19 annotated to the alpha subunit and 14 annotated to the beta subunit (Supplementary Information). Bayesian models were used to correlate Na^+^/K^+^-ATPase activity to transcript abundance. By comparing expected log pointwise predictive densities across models, we found mRNA abundance of beta subunits tended to have higher predictive power for enzyme activity than alpha subunits (Fig. 4).

### Behaviour

Alarm cue responses did not differ among experimental groups, both within and between the 0 h and 24 h timepoints used (Supplementary Fig. S7). No differences were observed among any rearing groups or trial groups combination, and no effect of cue was observed in time near the novel object (Supplementary Information). However, the group reared in high temperature but trialed in control conditions spent the highest proportion of time in the thigmotaxis zone, while the group reared in elevated *p*CO_2_ but trialed in control conditions trended toward less time in the thigmotaxis zone (Supplementary Information). There was no effect of food or alarm cue on time in the thigmotaxis zone.

With respect to total activity, between 50 and 53 days post hatch, the groups reared in elevated temperature and the combination of temperature and *p*CO_2_ exhibited greater total activity than the others, while the control and elevated *p*CO_2_-reared fish trialed in elevated *p*CO_2_ conditions trended toward less total activity, especially when exposed to an alarm cue (Fig. 5). There was no effect of alarm cue on total activity overall. No differences were observed in total activity near the novel object among rearing groups, trial groups, or cues (Supplementary Figs. S8-S11). Activity in the thigmotaxis zone was similar to total activity, with the greatest activity in the fish reared in elevated temperature but trialed in control conditions and a limited effect of cue (Supplementary Fig. S10). Overall, the cue delivered had limited or no effect on fish responses, while acute *p*CO_2_ exposure may have decreased activity. This effect of decreased activity from *p*CO_2_ exposure may have been masked by temperature which likely increased activity, especially in the combined *p*CO_2_ and temperature trial groups.

**Figure 5.**
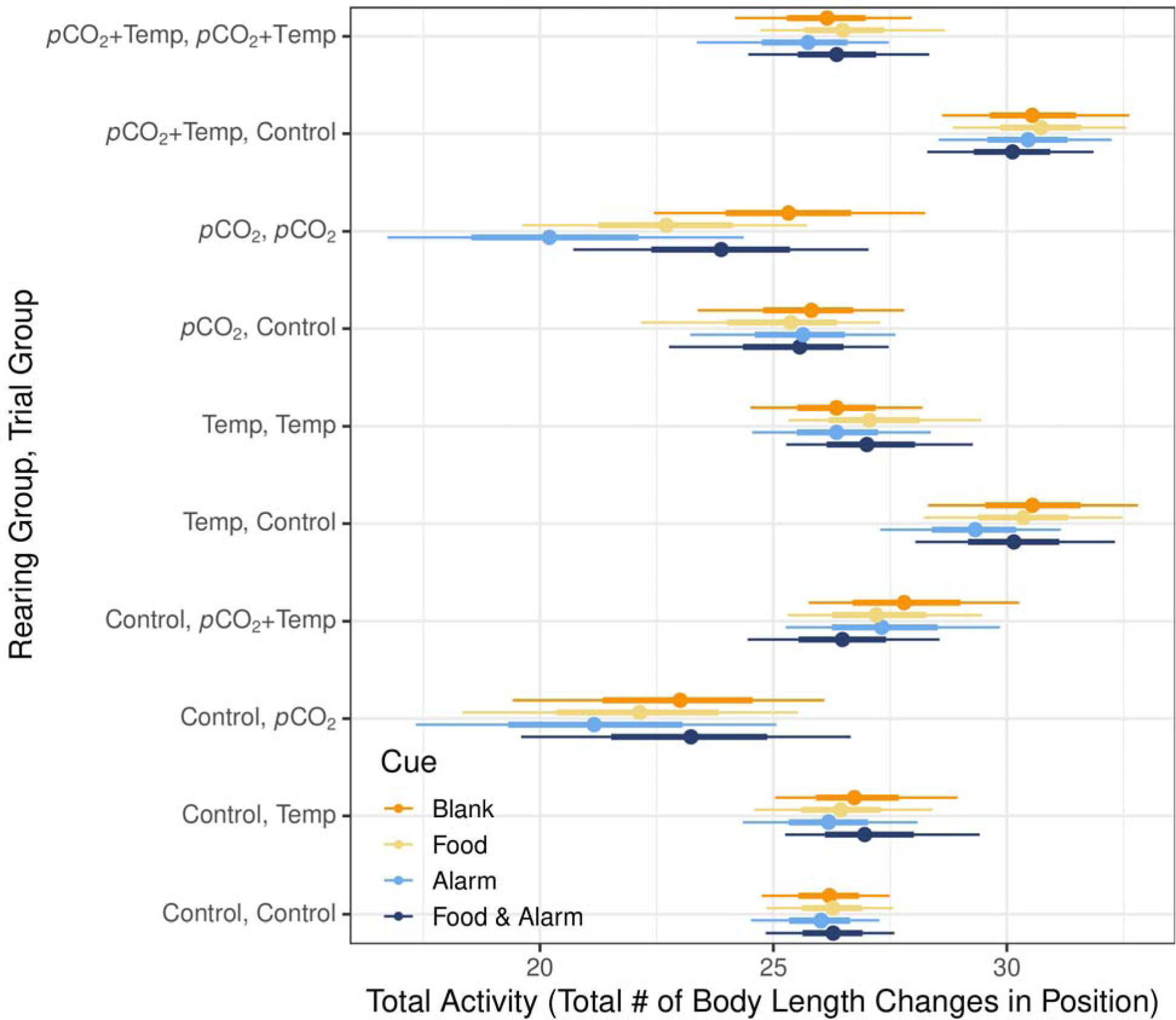
Credible intervals of total activity in crossed rearing and trial groups with respect to cue. Thin lines represent 95% credible intervals, while bold lines represent 66% credible intervals. Recordings were done for 5 minutes in the trial group conditions, with the middle 3 minutes of video used for analysis. Total activity was measured in total number of body length changes in position.

## Discussion

The present results revealed that increased developmental temperature and *p*CO_2_ independently as cumulative stressors influenced the responses to a transient 10,000 µatm *p*CO_2_ exposure in a simulated spring snowmelt. These findings hold several implications for freshwater fishes responding to climate change. Behaviourally, activity decreased in response to acute *p*CO_2_ exposure, regardless of rearing condition. Fish reared in elevated *p*CO_2_ and temperature exhibited transcriptional changes consistent with a decrease in organ growth, a pattern associated with lowered growth in response to CO_2_ in other freshwater species^1^. One gene potentially involved in the decrease in organ growth, sodium-dependent serotonin transporter, was involved in blood flow regulation and oxygen chemoreception in gill neuroephithelial cells^34,35^. In addition, compensatory mechanisms potentially consistent with a lowered routine metabolic rate were observed in the combined group. The transcriptional signal of a decrease in organ growth was consistent with the lower routine metabolic rate in fish reared in elevated *p*CO_2_ and temperature compared to a control group, 720 hours after the transient *p*CO_2_ exposure. The present work demonstrates that physiological and transcriptional processes remain different for at least 7 days post-exposure to transient *p*CO_2_ increase. Therefore, spring snowmelt with increasing atmospheric CO_2_ may have more pronounced effects on fish in watersheds characterized by heavy flows from snowpacks, depending on local geochemistry^36^. Thus, future research on the effects of climate change on seasonal acclimation in aquatic freshwater organisms should integrate the effects of CO_2_ and temperature over extended periods, as the factors together induce unique and prolonged physiological responses. Last, *p*CO_2_ will need to be considered in conservation management as it can induce responses independent from temperature, despite the two being tightly linked environmental variables.

Temperature alone exhibited the transcriptional network with highest connectivity based on betweenness scores, while the combination of both *p*CO_2_ and temperature had the least connectivity. Shared regulatory mechanisms were at the center of each network among the three treatments. However, differences in the outcomes of responses were apparent by analyzing the edges of networks, which are likely related to compensatory mechanisms. For example, otolith composition was affected by temperature but not *p*CO_2_ in a study using the same experimental design^32^. Similarly, both the control and temperature groups had increased Na^+^/K^+^-ATPase activity following the transient *p*CO_2_ increase, but not the two groups reared in elevated *p*CO_2_ conditions. This result suggests that compensatory mechanisms for responding to acute *p*CO_2_ may have been present in the groups that developed in increased *p*CO_2_, but not the other individuals. Based on models of transcript abundance and enzyme activity, beta subunits tended to be most predictive of Na^+^/K^+^-ATPase enzyme activity. Therefore, in the context of elevated temperature, Na^+^/K^+^-ATPase activity might be regulated by enzyme movement to cell membranes rather then differential production of the binding site, because of the beta subunit’s role in membrane binding^37^.

Food and alarm cue responses did not differ among rearing or trial conditions, while total activity was lower in all groups exposed to acute *p*CO_2_. Interestingly, the total activity of control and *p*CO_2_ rearing conditions were not different from each other following acute *p*CO_2_ exposure. Therefore, the exposure to *p*CO_2_ itself rather than developmental environment led to the observed differences. Elevated temperature could have mitigated the effects of *p*CO_2_ on total activity despite reduced olfactory responses to acidification^1,38–40^. Therefore, periods of high transient *p*CO_2_ but low temperature could influence organism’s energy budgets, with the potential for negative impacts on foraging and growth^41,42^. Marine acidification likely has inconsistent effects on fish behaviour^43^, and our results support those observations in freshwater fish.

Variability of *p*CO_2_ in flowing freshwater over time can be more than an order of magnitude greater than in marine systems^11,17,44–46^. While researchers may expect adaptations to CO_2_ flucuations in freshwater organisms^2,3,7^, this variability has nevertheless been shown to induce physiological responses in the lake sturgeon and other species^1^. Lake sturgeon, like other sturgeons, exhibit exceptional phenotypic plasticity to various environmental stressors^47–49^. This high plasticity makes sturgeons valuable for studying responses to environmental change at multiple biological scales, while their status as an ancient fish enables analyses of potentially conserved traits^50^. We observed extensive transcriptional and physiological plasticity in response to ecologically relevant changes in *p*CO_2_ and temperature. While other freshwater organisms may not have the capacity to marshal such a plastic response, elements of the cellular stress response are deeply conserved across all organisms^51,52^. Therefore, these may be critical response mechanisms for freshwater organisms faced with climate change.

## Acknowledgements

We thank Lindsey Belding, who provided access to necessary data under difficult circumstances. Joshua Sutherby assisted with lab work, and Alicia Korpach reviewed Bayesian analyses. Alyssa Weinrauch and Ian Bouyoucos engaged in helpful discussions about Na^+^/K^+^-ATPase data. This work was carried out on the original lands of the Anishinaabeg, Cree, Oji-Cree, Dakota and Dene peoples, and on the homeland of the Métis Nation. Water was sourced from the Shoal Lake 40 First Nation. Many bioinformatic steps in this research were enabled by our chance to use computing clusters of Prairies DRI and the Digital Research Alliance of Canada. Funding was provided by an NSERC/Manitoba Hydro Industrial Research Chair awarded to W.G.A., and Natural Science and Research Council of Canada Discovery Grants awarded to W.G.A. and K.M.J. (05348 and 05479, respectively).

## Data Availability

Raw messenger RNA sequencing data are available at the NCBI Sequence Read Archive (SRA Accession: PRJNA953937). Raw behavioural data tables and representative video files are available on Fig.share (https://doi.org/10.6084/m9.Fig.share.24043269.v1). All bioinformatic and statistical code used for the present analyses are available and annotated on GitHub (https://github.com/BioMatt/lkst_pCO2).

