## Supplementary Fig S. for "Integrated organismal responses induced by ecologically-relevant *p*CO_2_ and temperature exposures in developing lake sturgeon"

Supplementary Figure 1. Heatmap of log_2_ counts per million from RNAseq data for lake sturgeon of the different experimental groups, at different timepoints following a 10,000 µatm transient *p*CO_2_ increase.

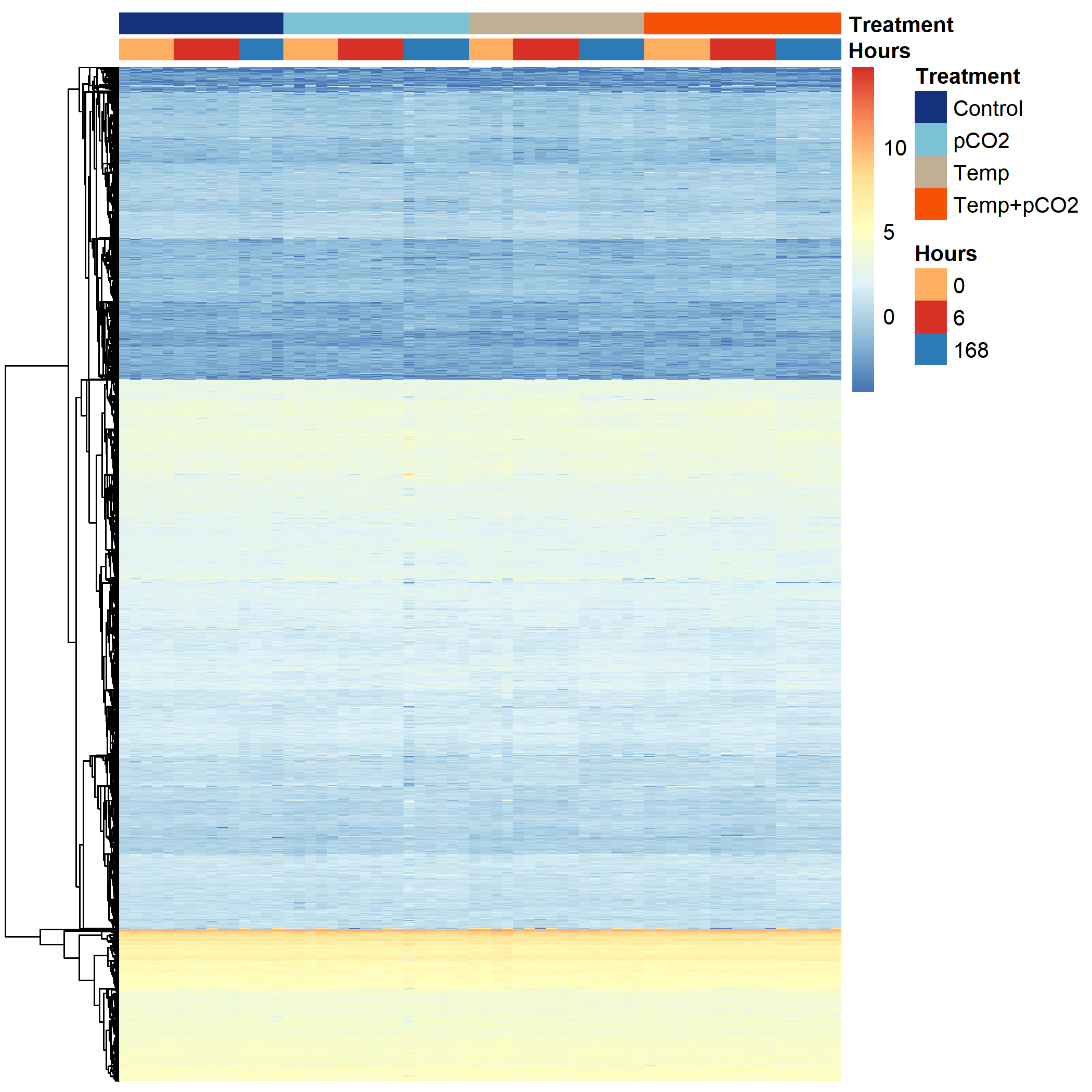

Supplementary Figure 2. Violin plots of messenger RNA transcript abundance of *slc6a4* in counts per million, separated into experimental groups and timepoints following a 10,000 µatm transient *p*CO_2_ increase.

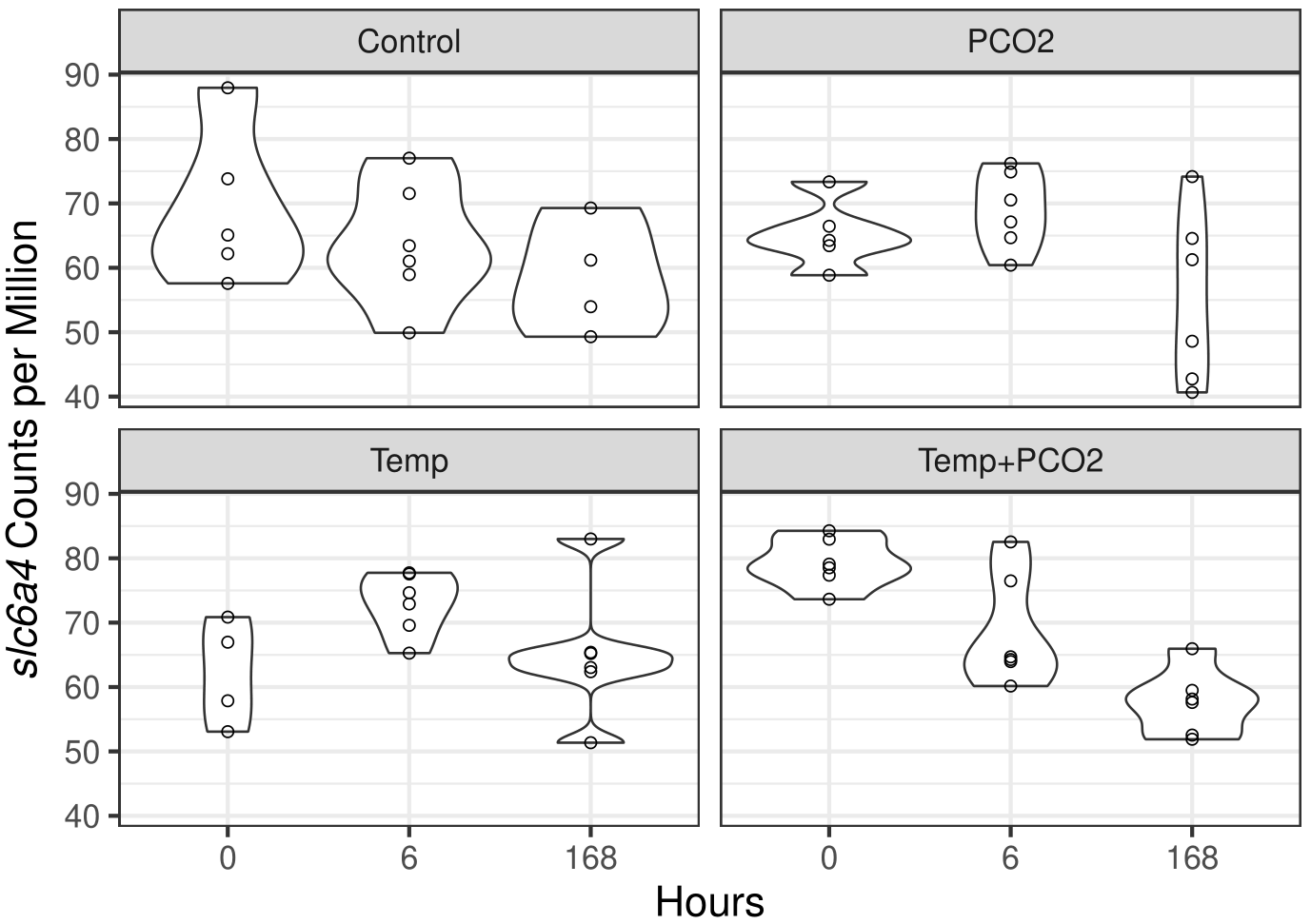

Supplementary Figure 3. Posterior distributions, 95% credible intervals (thin lines) and 66% credible intervals (thick lines) of hematocrit in lake sturgeon of different experimental groups and timepoints following a 10,000 µatm transient *p*CO_2_ increase.
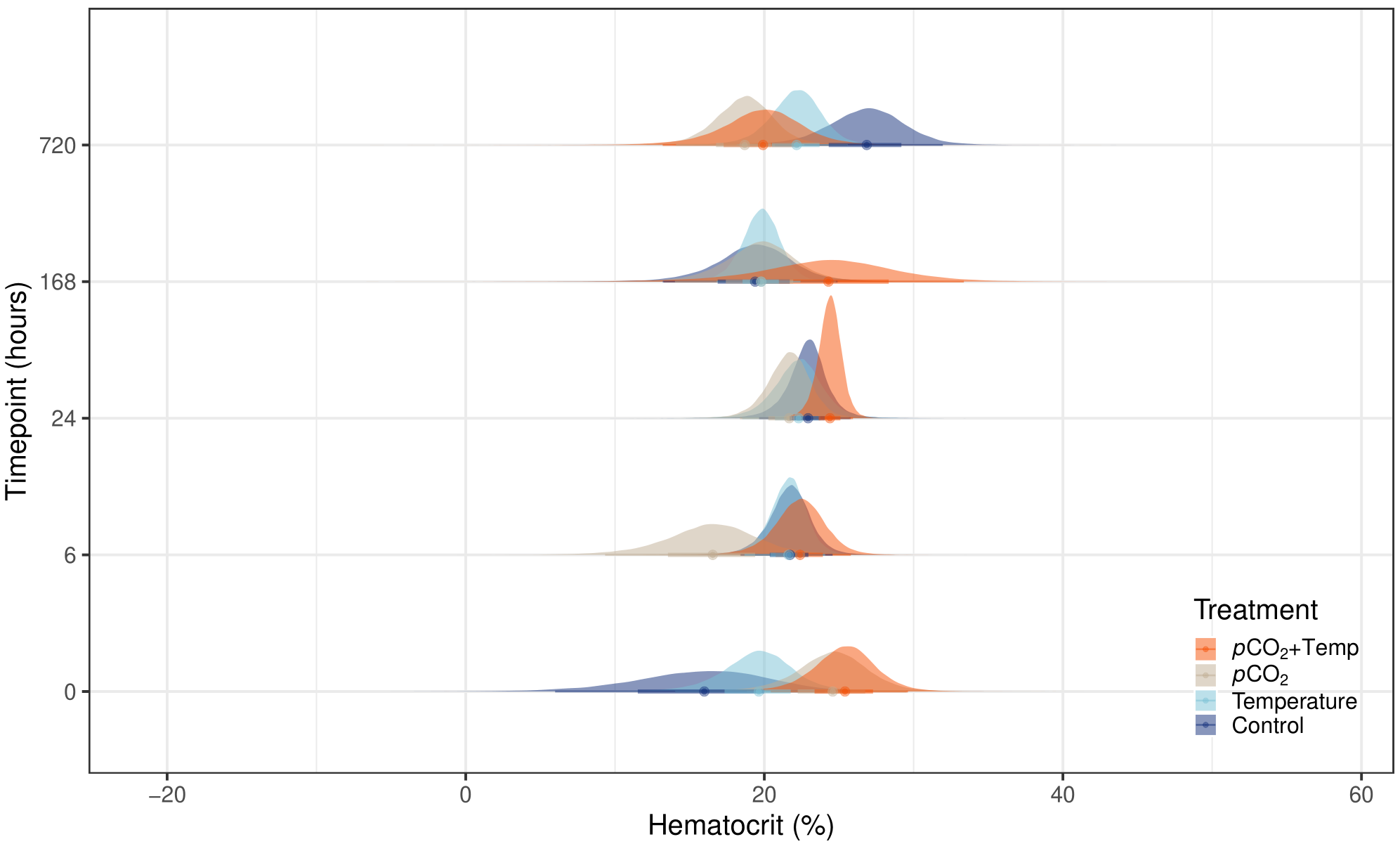

Supplementary Figure 4. Posterior distributions, 95% credible intervals (thin lines) and 66% credible intervals (thick lines) of ammonia excretion in lake sturgeon of different experimental groups and timepoints following a 10,000 µatm transient *p*CO_2_ increase.

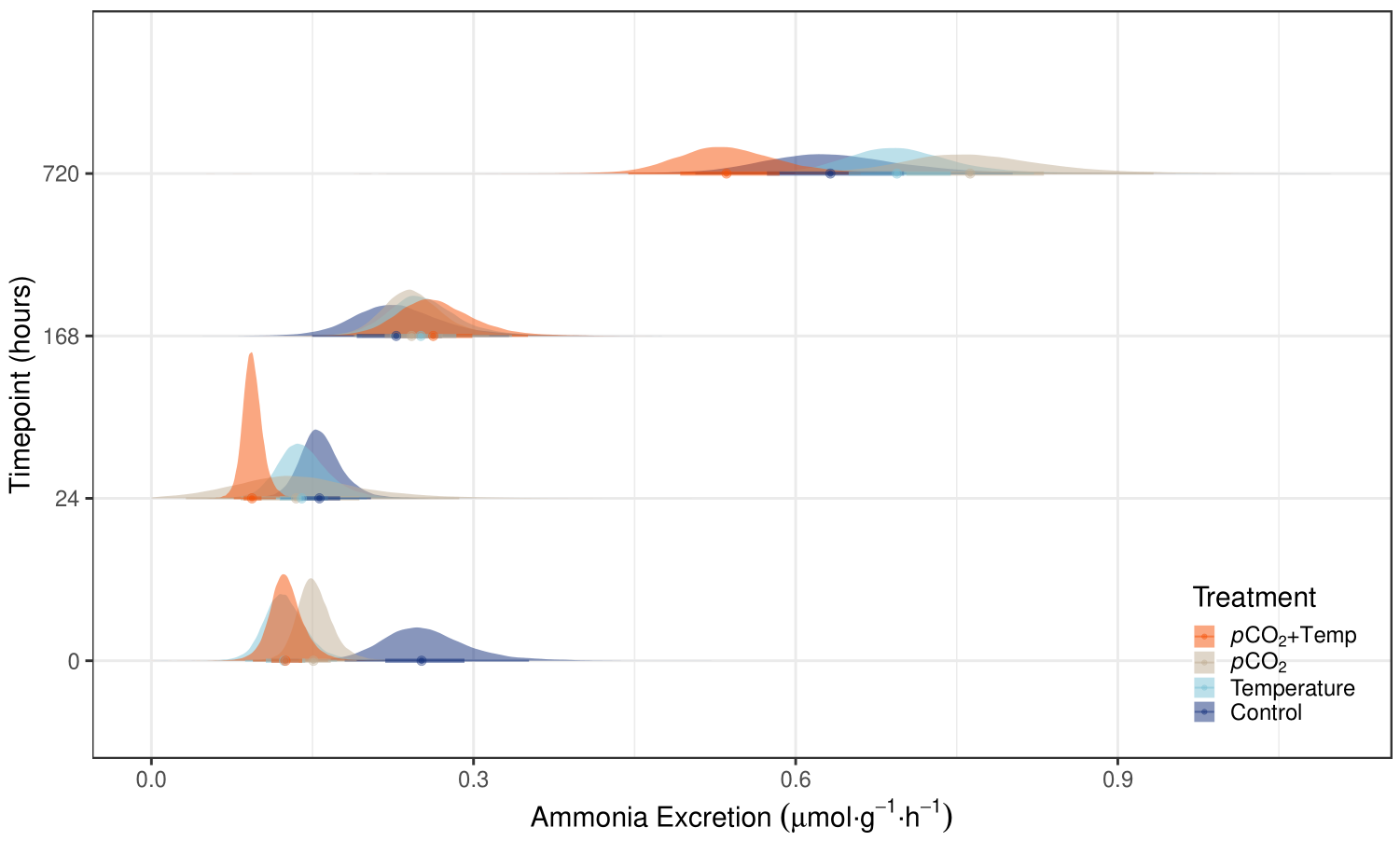

Supplementary Figure 5. Posterior distributions, 95% credible intervals (thin lines) and 66% credible intervals (thick lines) of A) routine and B) maximum metabolic rate in lake sturgeon of different experimental groups at 720h after the start of the 10,000 µatm transient *p*CO_2_ increase.

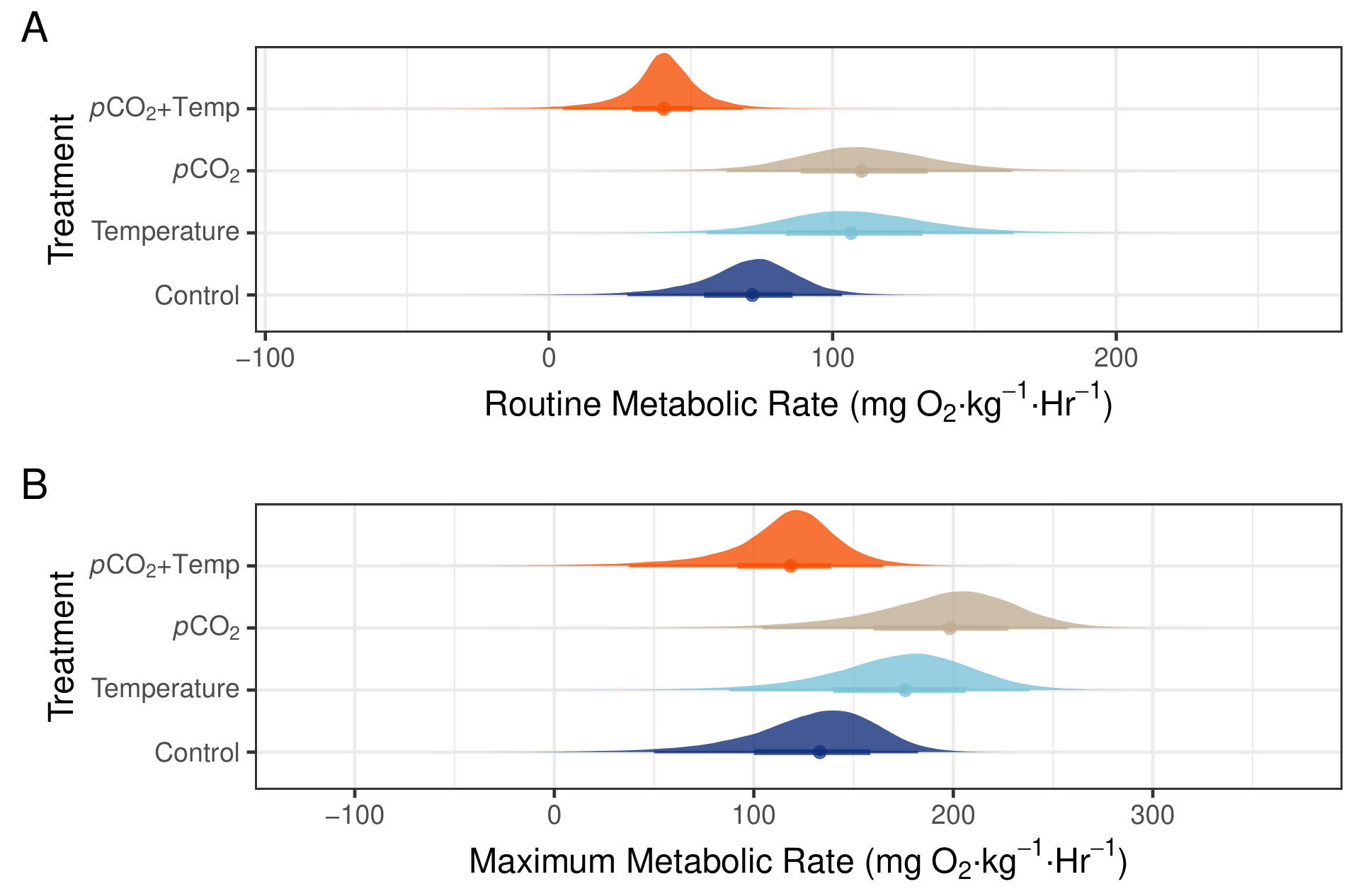

Supplementary Figure 7. Posterior distributions, 95% credible intervals (thin lines) and 66% credible intervals (thick lines) of proportion of time spent in alarm cues in lake sturgeon of different experimental groups and timepoints following a 10,000 µatm transient *p*CO_2_ increase.
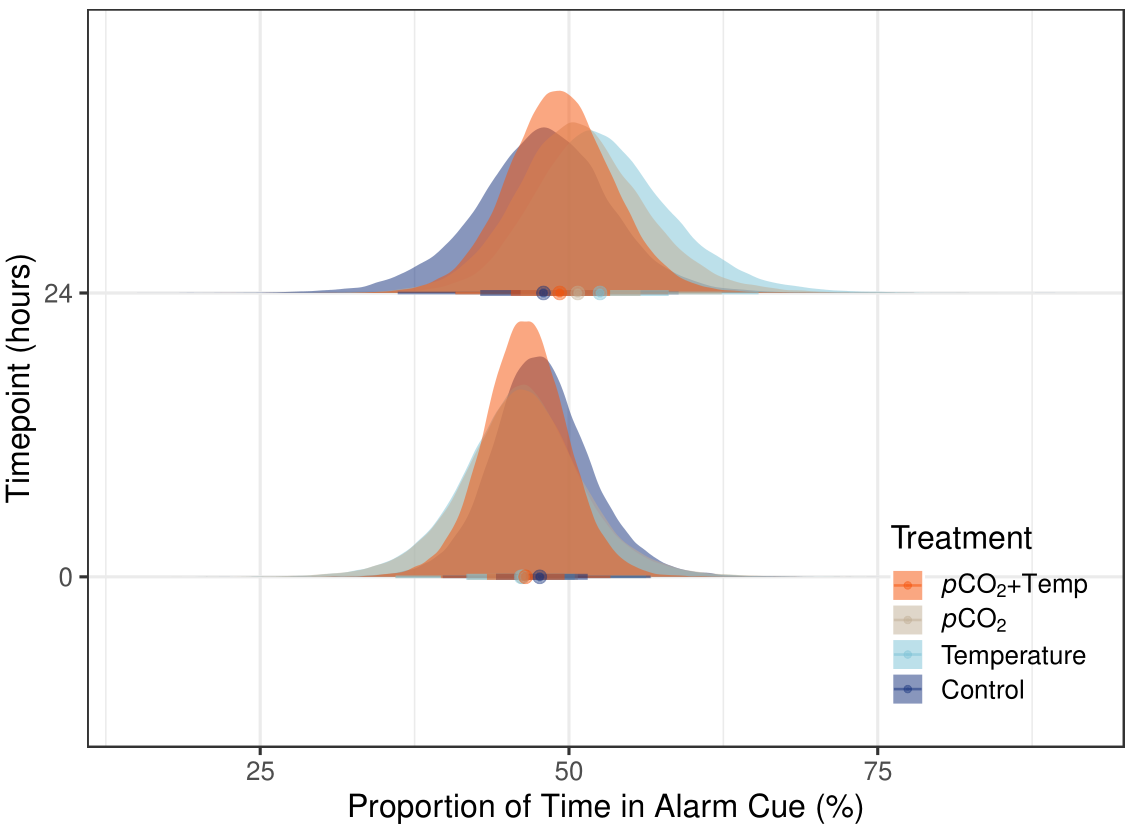

Supplementary Figure 8. 95% credible intervals (thin lines) and 66% credible intervals (thick lines) of total activity near a novel object in lake sturgeon of different rearing and experimental groups exposed to different cues.

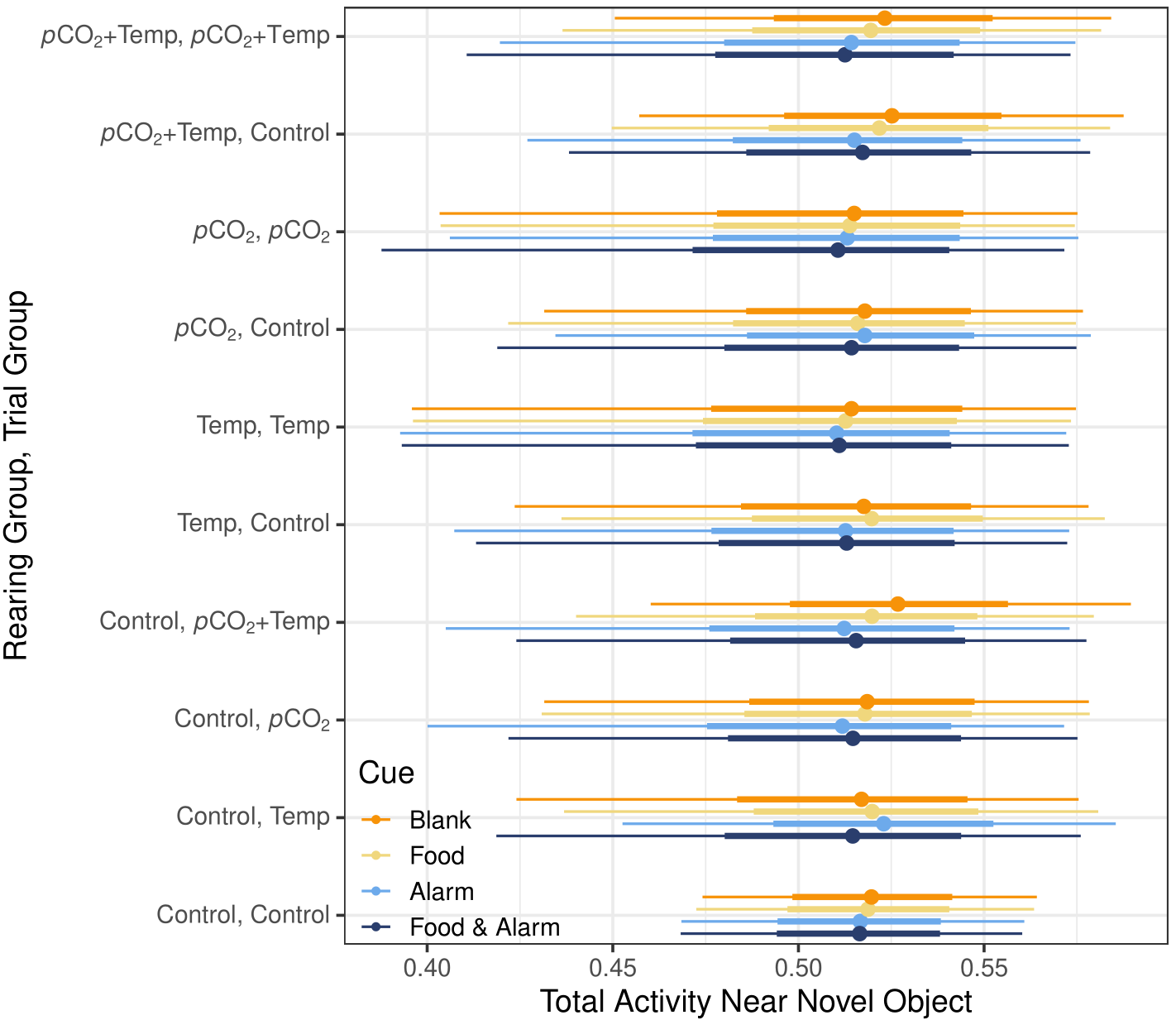

Supplementary Figure 9. 95% credible intervals (thin lines) and 66% credible intervals (thick lines) of proportion of time spent near a novel object in lake sturgeon of different rearing and experimental groups exposed to different cues.

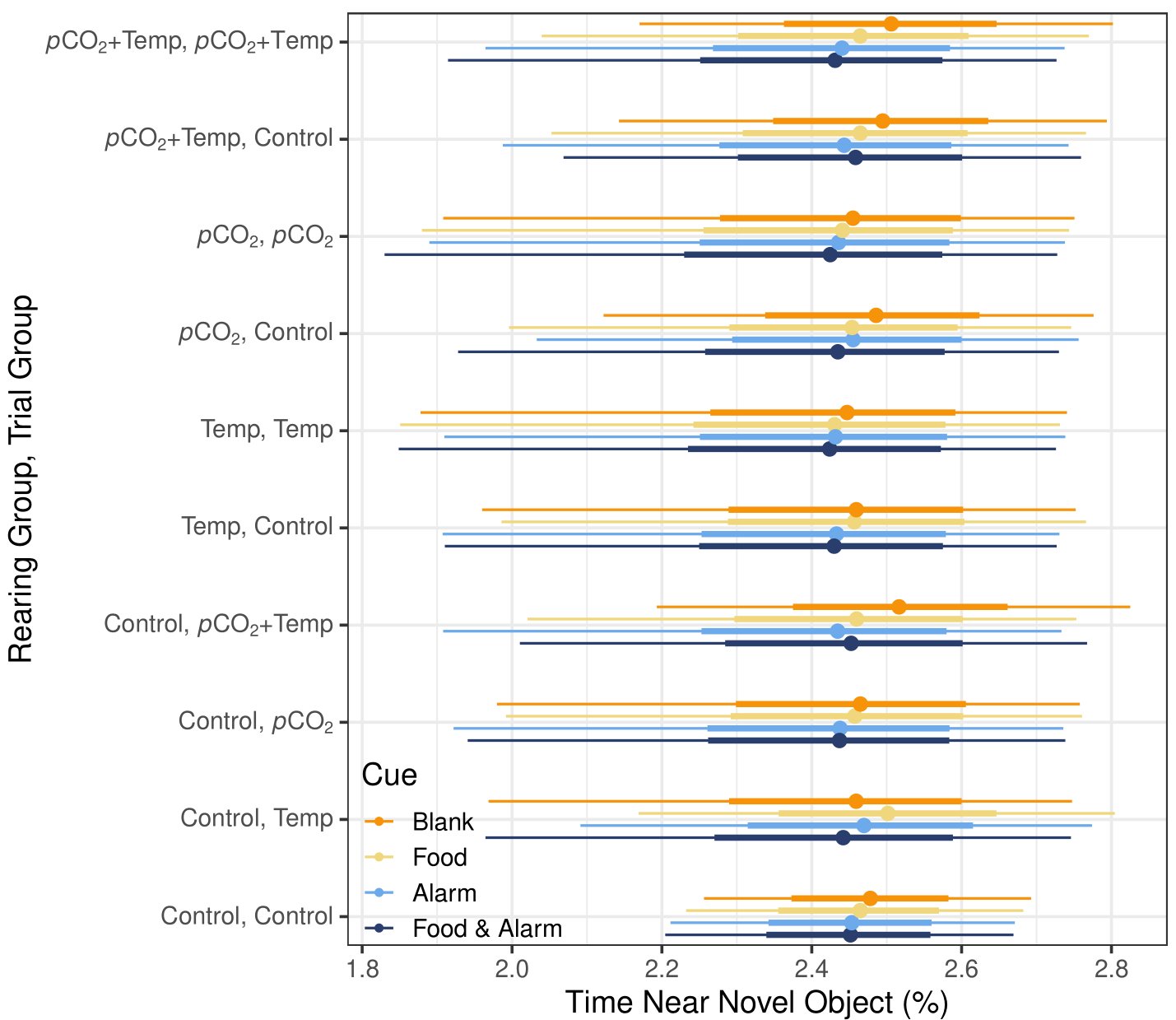

Supplementary Figure 10. 95% credible intervals (thin lines) and 66% credible intervals (thick lines) of total activity in the thigmotaxis zone in lake sturgeon of different rearing and experimental groups exposed to different cues.

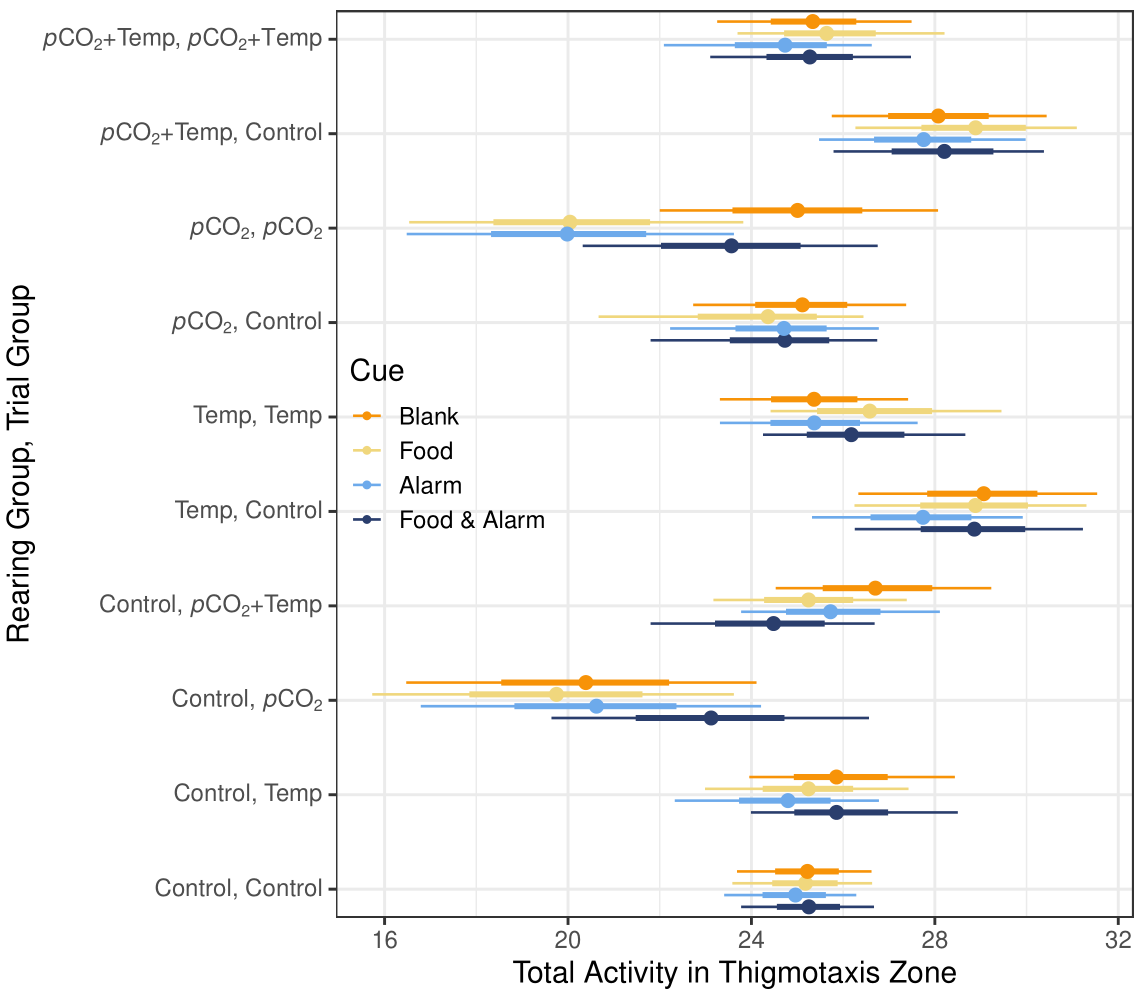

Supplementary Figure 11. 95% credible intervals (thin lines) and 66% credible intervals (thick lines) of proportion of time spent in the thigmotaxis zone in lake sturgeon of different rearing and experimental groups exposed to different cues.

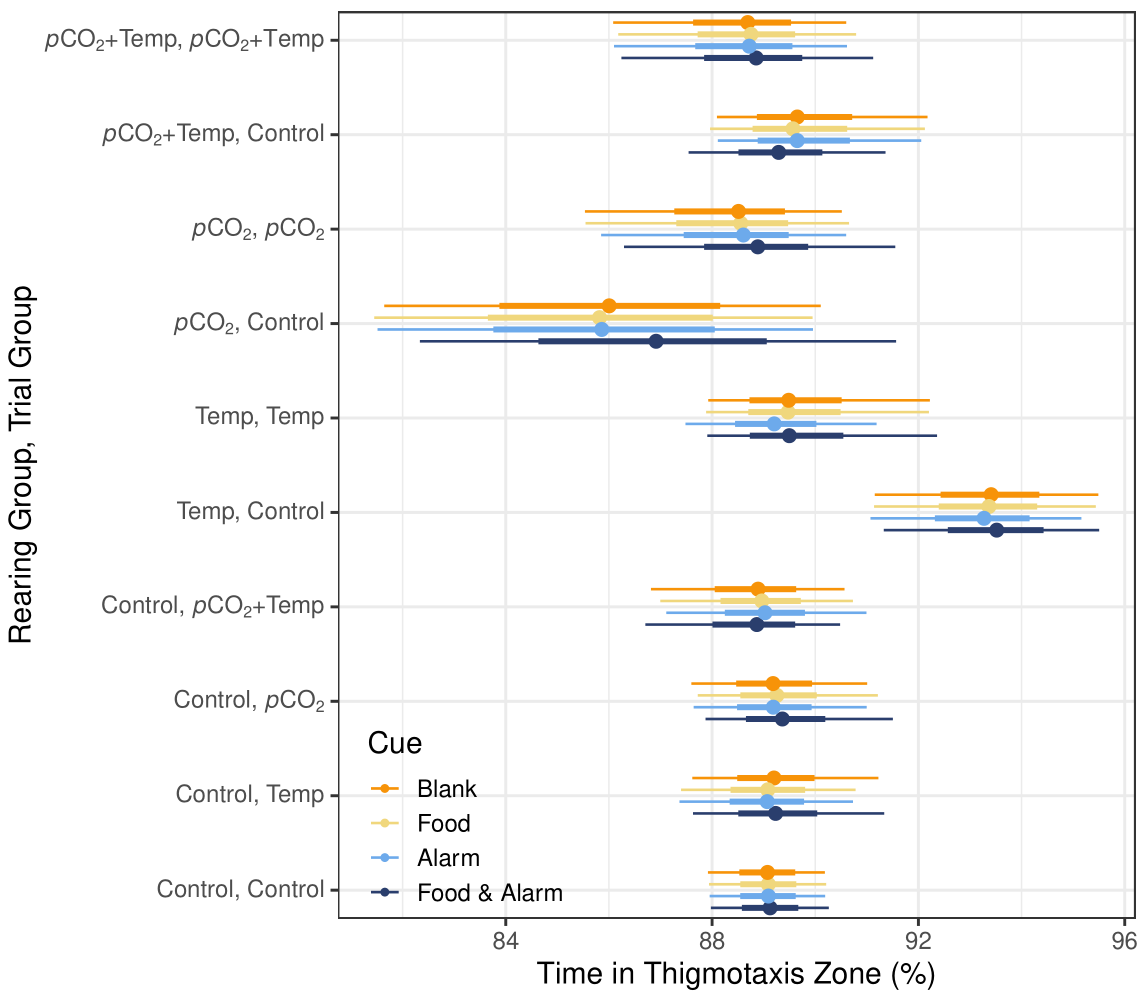
